# A chemically modified promoter-targeted duplex RNA activates *TCF4* expression

**DOI:** 10.64898/2026.10.02.756310

**Authors:** Bethany A. Janowski, Victor Tse, Jiaxin Hu, Sarah A. Cobb, Khayden Dong, David R. Corey

## Abstract

Pitt-Hopkins syndrome (PTHS) is a severe neurodevelopmental disorder caused by haploinsufficient expression of transcription factor 4 (*TCF4*). Increasing expression from the remaining functional *TCF4* allele could provide a mutation-independent strategy for patients with diverse pathogenic variants. We integrated genomic datasets with strand-specific RNA analysis to identify transcribed regions of the *TCF4* promoter suitable for targeting by activating double-stranded RNAs (dsRNAs). Screening of overlapping promoter-targeted dsRNAs identified a hotspot for increased *TCF4* expression. Activation was retained by a lead dsRNA containing extensive 2′-fluoro, 2′-O-methyl, and phosphorothioate modifications used in therapeutic oligonucleotides. Anti-AGO2 immunoprecipitation showed association with RNA transcribed from the targeted promoter region. dsRNA-mediated activation was associated with increased RNA polymerase II occupancy. These findings establish promoter-targeted dsRNAs as chemically tractable starting points for increasing *TCF4* expression and support their investigation as a mutation-independent strategy for haploinsufficient disease.

## INTRODUCTION

Short interfering RNAs (siRNAs) and antisense oligonucleotides (ASOs) are established drug development modalities for controlling gene expression (Egli and Manoharan 2023; Hofman and Corey 2024). Most approved siRNA and ASO drugs act by reducing the expression of disease-associated transcripts. ASOs can also increase therapeutic protein production by modulating splicing of a specific exon to compensate for a pathogenic gene variant. For example, the ASO nusinersen (Spinraza) enhances exon inclusion to increase levels of functional survival motor neuron (SMN) protein (Mercuri et al. 2018), while several exon-skipping ASOs increase production of a partially functional dystrophin as a treatment for Duchenne muscular dystrophy (Frank et al. 2020). These successes demonstrate the value of using oligonucleotides to increase production of proteins, but such strategies are limited to specific circumstances where changing splicing can increase expression of a therapeutic splice variant.

Haploinsufficient diseases are genetic conditions in which one allele of a gene is mutated, resulting in inadequate protein production (Johnson et al. 2019; Evans et al. 2024; Tse and Corey 2026). Haploinsufficiency is an important mechanism for genetic disease, with clinical evidence supporting involvement in hundreds of different diseases. In many cases, the mutations that cause a haploinsufficiency differ from patient to patient. This diversity complicates therapeutic development because effort would need to be divided among mutation-specific strategies. By contrast, an approach that increases expression of the wild-type gene would have the potential to treat many patients whose disease is caused by insufficient expression.

Promoter-targeted dsRNAs are a potentially general strategy for increasing the expression of endogenous wild-type proteins (Li et al. 2006; Johnson and Corey 2023). Our laboratory and others have observed that synthetic double-stranded RNAs (dsRNAs) that target gene promoters can be used to inhibit or activate gene expression (Kim et al. 2006; Li et al. 2006; Janowski et al. 2007; Place et al. 2008; Schwartz et al. 2008; Chu et al. 2010; Matsui et al. 2010; Matsui et al. 2013; Portnoy et al. 2016; Weinberg and Morris 2016; Matsui and Corey 2017; Reebye et al. 2018; Sarshad et al. 2018; Sarker et al. 2020; Johnson and Corey 2023; Bi et al. 2024; Desaulniers et al. 2025; Musiała-Kierklo et al. 2025; Qian et al. 2025; Shcherbinina et al. 2026). These promoter-targeted dsRNAs are designed using chemical architectures like conventional siRNAs and act in association with Argonaute (AGO) proteins (Kim et al. 2006; Li et al. 2006; Chu et al. 2010; Matsui et al. 2010; Matsui et al. 2013; Portnoy et al. 2016; Sarshad et al. 2018; Johnson and Corey 2023). The guide RNA strand (gRNA) provides sequence-specific recognition, whereas AGO protects the gRNA and facilitates binding to target sequences. Previous studies indicate that promoter-targeted dsRNAs can recognize RNA transcripts that overlap gene promoters, acting *in cis* relative to chromosomal DNA (Schwartz et al. 2008; Matsui et al. 2010; Matsui et al. 2013; Portnoy et al. 2016; Johnson and Corey 2023). Recognition of these promoter-associated RNAs by AGO:gRNA complexes can alter transcription at the corresponding genomic locus. We have found that dsRNAs containing central mismatches are particularly useful for this application because they can promote target recognition while reducing the potential for AGO2-mediated target cleavage of complementary sequences (Wang et al. 2008; Hu et al. 2010). Promoter-targeted dsRNAs can increase RNA polymerase II (RNAP2) occupancy at gene promoters and increase levels of gene transcription. These observations suggest that synthetic dsRNAs, in combination with AGO, can act as programmable regulators of gene expression.

In this paper, we use promoter-targeted dsRNAs to increase expression of *TCF4*, the gene whose mutation is responsible for the haploinsufficiency that causes Pitt-Hopkins Syndrome (PTHS), a severe neurodevelopmental disorder (Amiel et al. 2007; Brockschmidt et al. 2007; Zweier et al. 2007; Whalen et al. 2012; Zollino et al. 2019; Chen et al. 2021; Sirp et al. 2021; Teixeira et al. 2021; Kim et al. 2022; Papes et al. 2022; James et al. 2025; Bourin 2026). Approximately 1,600 patients have been identified worldwide, and there is substantial unmet medical need. Because pathogenic variants are distributed throughout TCF4 (Whalen et al. 2012; Zollino et al. 2019; Papes et al. 2022), increasing expression from the remaining functional allele could provide a mutation-independent strategy applicable across diverse patient genotypes and address the underlying deficiency in TCF4 expression.

We characterize the transcriptional landscape at the *TCF4* promoter and use this information to design synthetic promoter-targeted dsRNAs. We identify several related dsRNAs that increase *TCF4* expression, with most activating dsRNAs clustered at a sequence-defined hotspot. *TCF4* activation is retained by a dsRNA that contains the extensive 2’-fluoro, 2’-O-methyl, and phosphorothioate modifications commonly found in therapeutic siRNAs (Egli and Manoharan 2023). AGO2 immunoprecipitation recovers RNA transcribed from the targeted promoter region, and RNAP2 chromatin immunoprecipitation indicates that dsRNA-mediated activation is associated with increased RNAP2 occupancy. These data demonstrate that promoter-targeted dsRNAs have the potential to provide chemically tractable starting points for increasing *TCF4* expression and support investigating promoter RNAs a mutation-independent strategy for haploinsufficient disease.

## RESULTS

### Characterizing the DNA and RNA landscape of TCF4

The mechanism we and others previously characterized for RNA-mediated transcriptional activation requires the existence of a transcript that overlaps the promoter (Schwartz et al. 2008; Matsui et al. 2010; Matsui et al. 2013; Portnoy et al. 2016; Johnson and Corey 2023). Our first experimental goal, therefore, was to identify transcribed regions at the *TCF4* promoter that might be targets for activating dsRNAs. We chose A549 cells for this analysis because they express detectable TCF4 protein and can be efficiently transfected. Possession of a straightforward expression assay facilitated screening multiple candidate dsRNAs.

The *TCF4* locus is controlled by a TATA-less promoter and has a complex landscape of overlapping transcripts. We analyzed GENCODE annotations and identified three mRNA transcripts expected to encode the full-length TCF4 protein (**Supplemental Figure S1A**) (Mudge et al. 2025). The most broadly expressed transcript, ENST00000354452.8, is designated as the Matched Annotation from NCBI and EMBL-EBI (MANE) Select transcript and we prioritized it as the principal clinically relevant mRNA isoform for our study (Wright et al. 2023).

To gain more precise insight into transcription mechanisms and start sites, we defined promoter regions by integrating datasets from the ENCODE (He et al. 2020), CpG island (Saxonov et al. 2006), EPDnew (Dreos et al. 2013), and FANTOM5 CAGE (FANTOM Consortium and the RIKEN PMI and CLST (DGT) 2014) databases (**Supplemental Figure S1B,C**). This analysis identified three promoter-like regions, designated TSS1, TSS2, and TSS3 (**Figure 1A**). With our focus on the MANE-select transcript, we found that TSS1 aligned to the most 5’ end of this prioritized mRNA isoform. and Mapped CAGE data, and coinciding promoter-associated features annotated by ENCODE, CpG island analysis, and EPDnew, suggested TSS1 to be the most prominent promoter region with an active transcription start site. The CAGE data also revealed antisense transcription across the *TCF4* promoter region, showing that both sense and antisense RNA species were present across TSS1. These observations are consistent with TSS1 as the highest-priority target for complementary synthetic dsRNAs.

**Figure 1.**
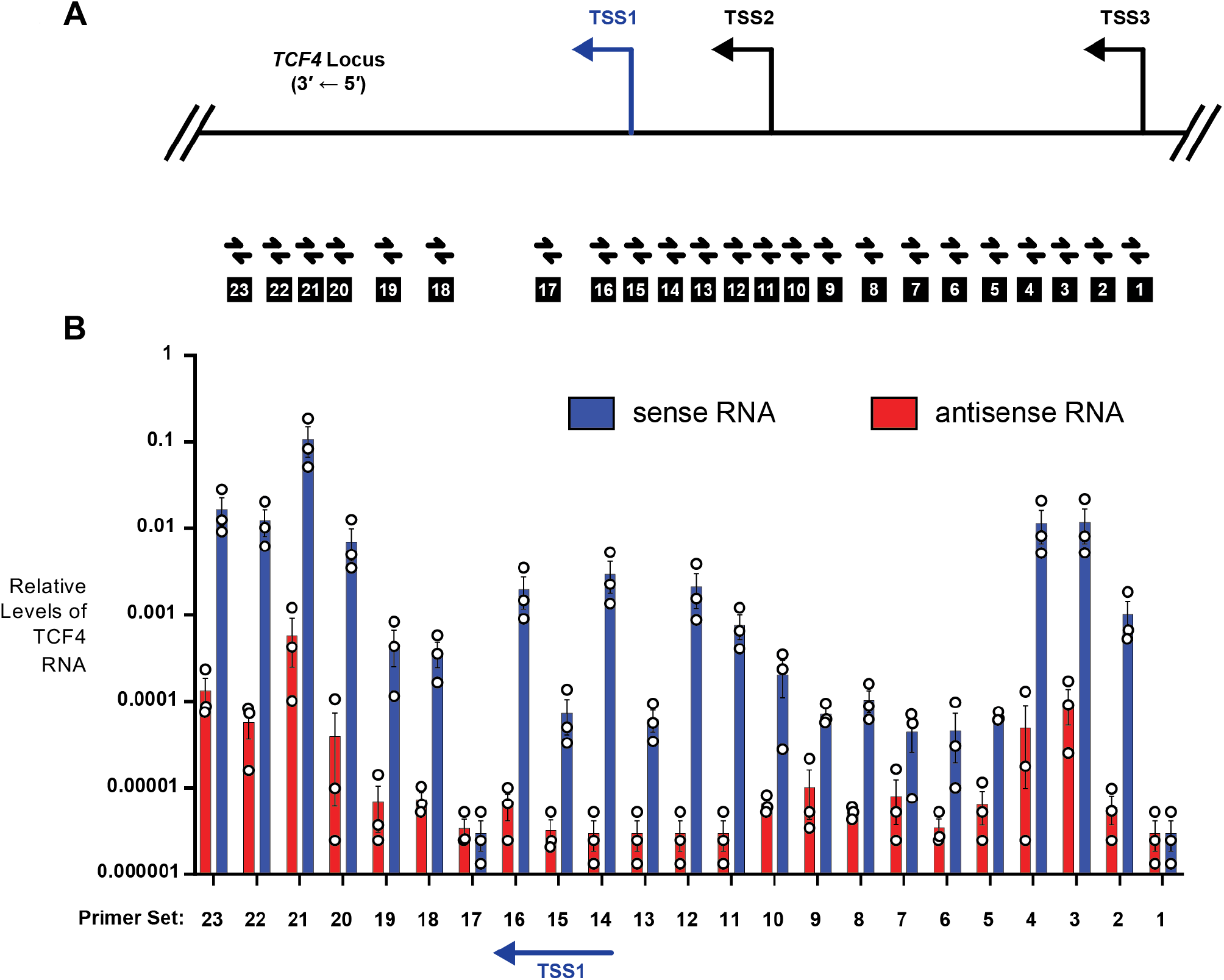
Transcription at the *TCF4* promoter. **(*A*)** Scheme showing TSS1, TSS2, and TSS3, the most prominent regions for initiating *TCF4* transcription. TSS1 is the primary target for this study. **(*B*)** Experimental characterization of transcription. Top: location of qPCR primer pairs relative to TSS1, TSS2, and TSS3. Bottom: qPCR (log scale) showing sense and antisense transcription across the *TCF4* promoter. Sense strand transcription is annotated in blue. Antisense strand transcription is indicated in red. Database analysis of the *TCF4* promoter is shown in Supplemental Figure S1. All data are from ≥ 3 independent biological replicates.

To confirm the existence of transcription upstream and downstream from TSS1, we used strand-specific reverse transcription followed by qPCR across the region surrounding TSS1 (**Figure 1B**) (Matsui et al. 2013). We detected both sense and antisense transcripts overlapping regions upstream and downstream of TSS1. We employed a separate qPCR experiment using cDNA templates generated from “plus reverse transcription” and “minus reverse transcription” reactions performed in parallel (**Supplemental Figure S1D**) to confirm that specificity in targeting and detecting only RNA species by this assay, and no genomic or contaminating DNA, These data are consistent with the genomic datasets analyzed and provide the information necessary to design potential activating dsRNAs.

### Design and screening of promoter-targeted dsRNAs

After characterizing transcription across the *TCF4* promoter, we designed dsRNAs complementary to regions adjacent to the major transcription start sites. The dsRNAs used a conventional siRNA-like architecture but contained a mismatch at position 9 of the guide strand relative to target. This mismatch permits target recognition while reducing the potential for AGO2-mediated cleavage (Wang et al. 2008; Hu et al. 2010). A second mismatch at position 19 of the passenger strand was included to favor loading of the intended guide strand onto AGO proteins (Schwarz et al. 2003).

Candidate dsRNAs were designed as an overlapping series in which adjacent compounds differed by one nucleotide. This tiling strategy allowed us to comprehensively test a region for its ability to support productive activation rather than relying on the activity of a single sequence. dsRNAs were introduced into A549 cells by lipid-mediated transfection, and endogenous *TCF4* expression was evaluated by qPCR and western blotting. Positive and negative controls followed standard design criteria (Gagnon and Corey 2019). Experiments included a non-complementary dsRNA, a scrambled dsRNA derived from an active sequence identified in our study, and a fully complementary siRNA directed against *TCF4* mRNA as a positive control for successful transfection and RNA interference activity.

Because compensating for haploinsufficiency may involve relatively small fold changes, and since we sought to quantify differences among multiple compounds using western blotting, we emphasized independent biological replication and appropriate statistical analysis (Gassmann et al. 2009; Taylor et al. 2013; Taylor and Posch 2014). For each condition tested, we obtained data from at least three independent biological replicates generated by separate transfections (**Supplemental Figure S3B**). Technical replicates were averaged within each independent biological replicate. We assessed the distribution and variance of western blot quantification data by statistical tests. These tests included quantile-quantile plots and a Brown-Forsythe test to establish whether the data met the assumptions needed to perform a statistical test.

### Tiled dsRNAs identify a promoter hotspot for *TCF4* activation

We prioritized the TSS1 promoter region because it showed the strongest initial evidence of promoter activity and transcription. dsRNAs TSS1-8 through TSS1-29 were designed to target TSS1, overlapping one another and differing by single-nucleotide increments (**Figure 2A**). Our initial screen identified TSS1-15 as a reproducible activator and this duplex was selected as the benchmark compound for subsequent experiments (n=28 independent replicates). A negative scrambled control based on the sequence of TSS1-15 (TSS1-15SCR) was used in all experiments. We used siRNA (E18) as a positive control for transfection efficiency. siRNA E18 is fully complementary to *TCF4* mRNA exon 18 (Forrest et al. 2013). Exon 18 serves as an ideal target for a positive control siRNA because it is shared by a majority of *TCF4* transcript variants. Application of E18 produced a lower molecular weight band of unknown origin which does not interfere with detecting gene knockdown of full-length TCF4 protein.

**Figure 2.**
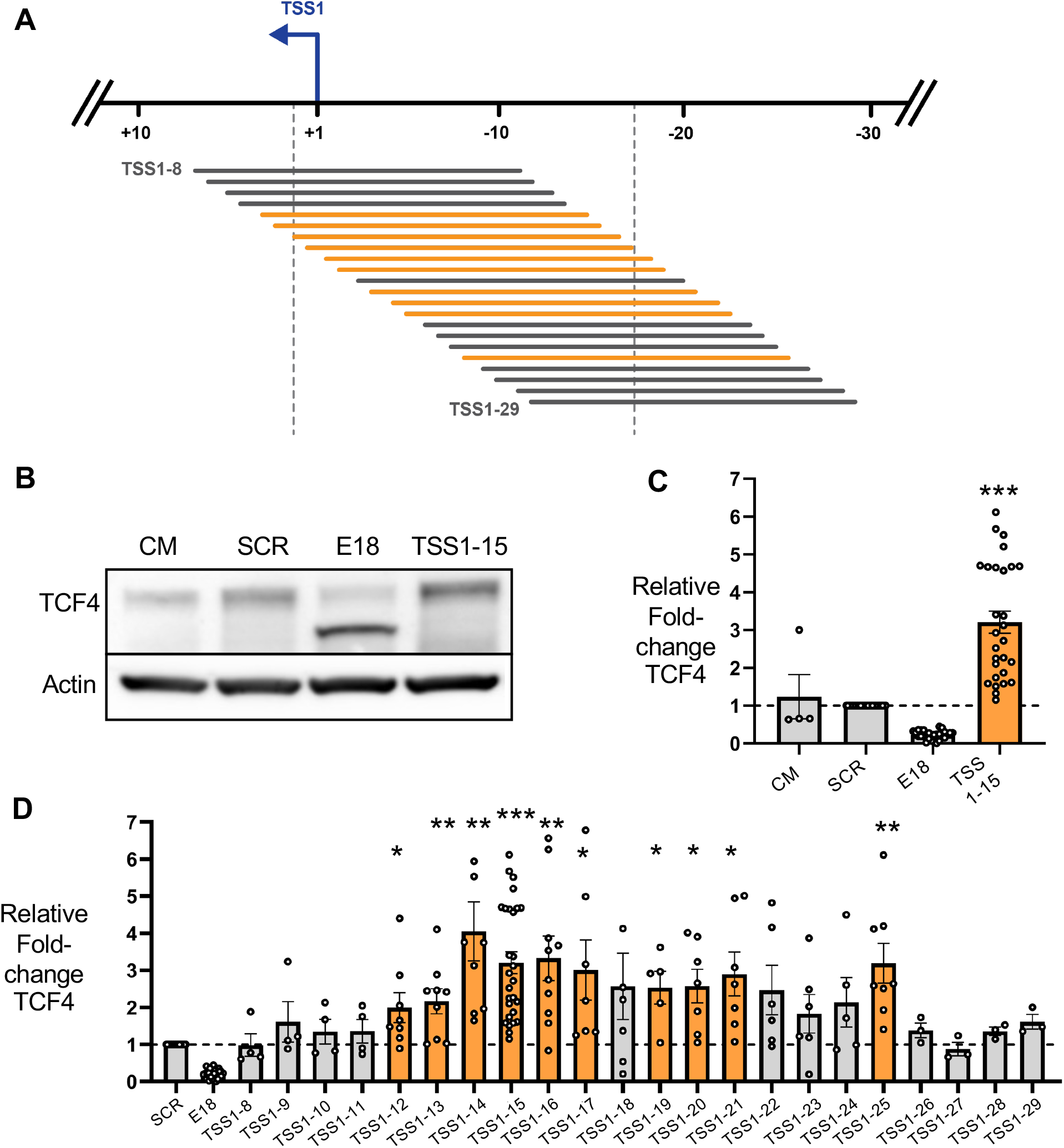
Activation of TCF4 protein expression by dsRNAs that target TSS1. **(*A*)** Schematic showing location of duplex RNAs TSS1-8 to TSS1-29. Promoter-targeted duplex RNAs that showed statistically significant activation >1.5 fold increased gene expression are orange. **(*B*)** Western analysis of activation by benchmark TSS1-15 relative to controls. **(*C*)** Quantification of all independent western analysis replicates of effect of TSS1-15 on TCF4 protein expression. **(*D*)** Quantification of western analyses of TCF4 protein expression after addition of TSS1-8 to TSS1-29. Promoter-targeted duplex RNAs that showed a statistically significant activation >1.5 fold are shown in orange. Sample significance was then determined using a two-tailed Welch’s t-test. Statistical significance is indicated as follows: ∗p ≤ 0.05, ∗∗p ≤ 0.01, and ∗∗∗p ≤ 0.001. n ≥ 3 independent experimental replicates, individual values are shown. Details of replicate number and p-values are described in Supplemental Figure 3. CM is a noncomplementary control duplex.

Duplex RNA TSS1-15 increased TCF4 protein expression by approximately 3-fold (**Figure 2B,C; Supplemental Figure S2A**). We observed statistically significant activation by RNAs that were targeted to adjacent or nearby sequences, including TSS1-12, TSS1-13, TSS1-14, TSS1-16, TSS1-17, TSS1-19, TSS1-20, TSS1-21, and TSS1-25 (**Figure 2D; Supplemental Figure S2B, Supplemental Figure S3B**). Taken together, these data identify a sequence-defined hotspot that supports productive activation. Because this activation is achieved by several related dsRNAs even though the seed sequences for recognition differ among the active compounds, our data support the suggestion that activation is due to on-target recognition rather than a seed-sequence dependent off-target effect.

### A fully chemically modified duplex retains *TCF4*-activating activity

Unmodified dsRNAs are rapidly degraded in biological systems and require chemical modification to be suitable starting points for *in vivo* development (Egli and Manoharan 2023; Hofman and Corey 2024). Chemical modification, however, can alter dsRNA recognition, AGO loading, and target engagement. To determine whether TCF4 activation was compatible with chemical architectures similar to those of conventional therapeutic duplex RNAs, we tested Mod15, an extensively modified analogue of TSS1-15. Mod15 contains 2′-fluoro and 2′-O-methyl nucleotides at every ribose position together with terminal phosphorothioate linkages (**Figure 3A**) (Egli and Manoharan 2023). There were no unmodified ribonucleotides. A noncomplementary duplex (Mod-Control) containing the same modification pattern was included as a negative control.

**Figure 3.**
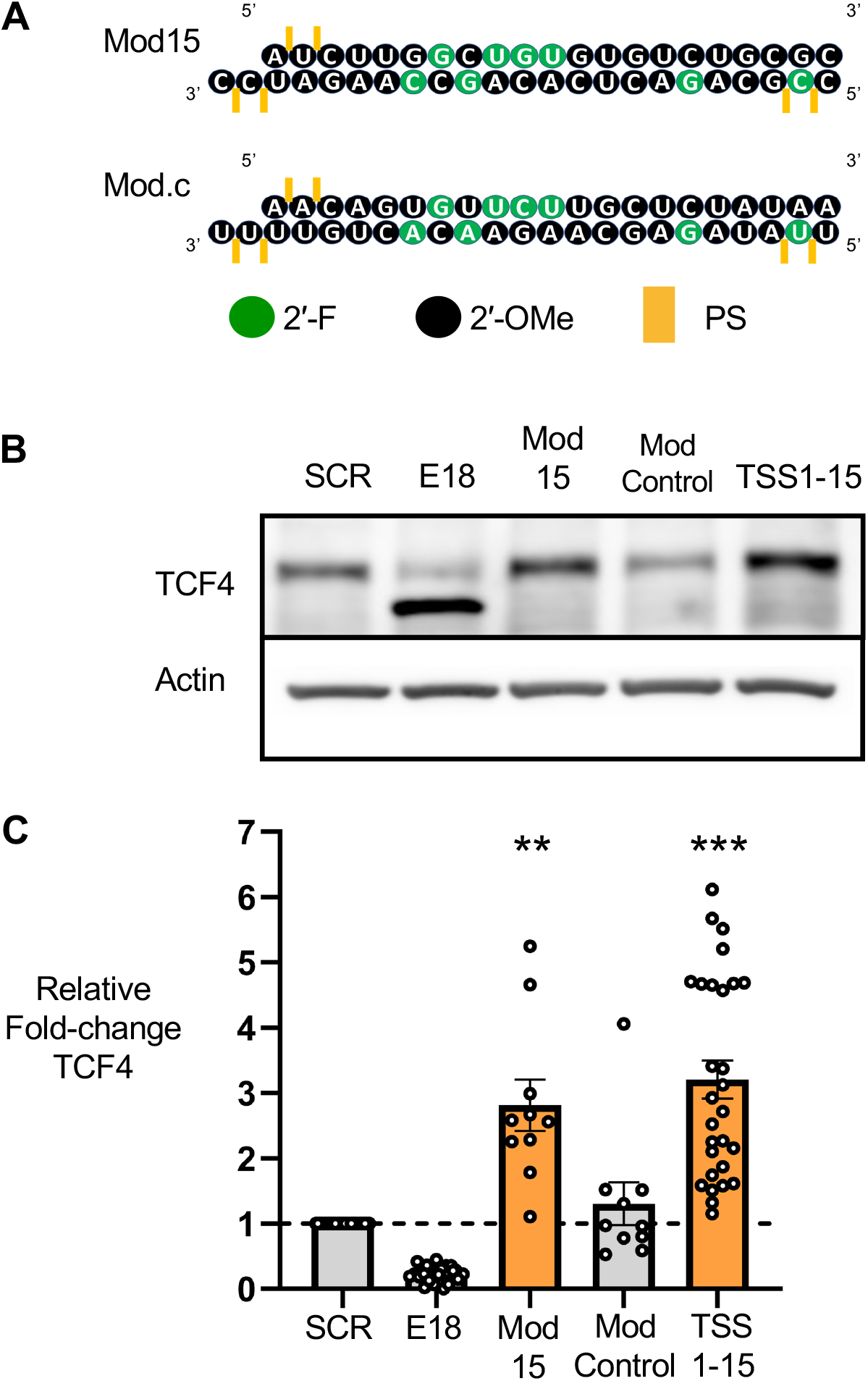
Activation of TCF4 protein expression by a chemically modified promoter-targeted RNA. **(*A*)** Sequences and chemical modifications of promoter-targeted RNA Mod15 and a noncomplementary control with an identical pattern of modifications. 2’-F = 2’-fluoro; 2’-Ome = 2’-O-methyl; PS = phosphorothioate. **(*B*)** Western analysis of TCF4 activation after addition of Mod15 relative to its corresponding control (Mod-Control), compared to its unmodified RNA analogue TSS1-15 compared to its corresponding control (SCR). **(*C*)** Quantification of (*B*). Sample significance was then determined using a two-tailed Welch’s t-test. Statistical significance is indicated as follows: ∗p ≤ 0.05, ∗∗p ≤ 0.01, and ∗∗∗p ≤ 0.001. Details of replicate number and p-values are described in Supplemental Figure S3.

Mod15 significantly increased TCF4 protein expression, whereas Mod-Control did not (**Figure 3B,C; Supplemental Figure S3**). The magnitude of activation by Mod15 was similar to activation by the unmodified parent dsRNA, TSS1-15. These results demonstrate that extensive chemical modification using chemistries commonly incorporated into therapeutic oligonucleotides is compatible with promoter-targeted activation of *TCF4*.

### AGO2 associates with RNA transcribed from the *TCF4* promoter region

Previous studies suggest that promoter-targeted dsRNAs act through AGO-containing complexes that recognize RNA transcripts overlapping gene promoters (Schwartz et al. 2008; Matsui et al. 2010; Matsui et al. 2013; Portnoy et al. 2016; Johnson and Corey 2023). To determine whether AGO2 associates with RNA expressed from the targeted *TCF4* promoter region, we performed RNA immunoprecipitation using an anti-AGO2 antibody after transfection with our lead activator TSS1-15.

We observed that promoter-region RNA was enriched in anti-AGO2 immunoprecipitates after transfection with TSS1-15 when compared to transfection with duplex SCR. We did not observe enrichment when control IgG was used (**Figure 4A; Supplemental Figure S4A**). These findings support a model in which AGO2-guide RNA complexes recognize RNA transcribed from the targeted promoter region, consistent with mechanisms previously described for promoter-targeted activation of the *LDL receptor* (Matsui et al. 2010), *progesterone receptor* (Schwartz et al. 2008), and *cyclooxygenase-2* genes (Matsui et al. 2013).

**Figure 4.**
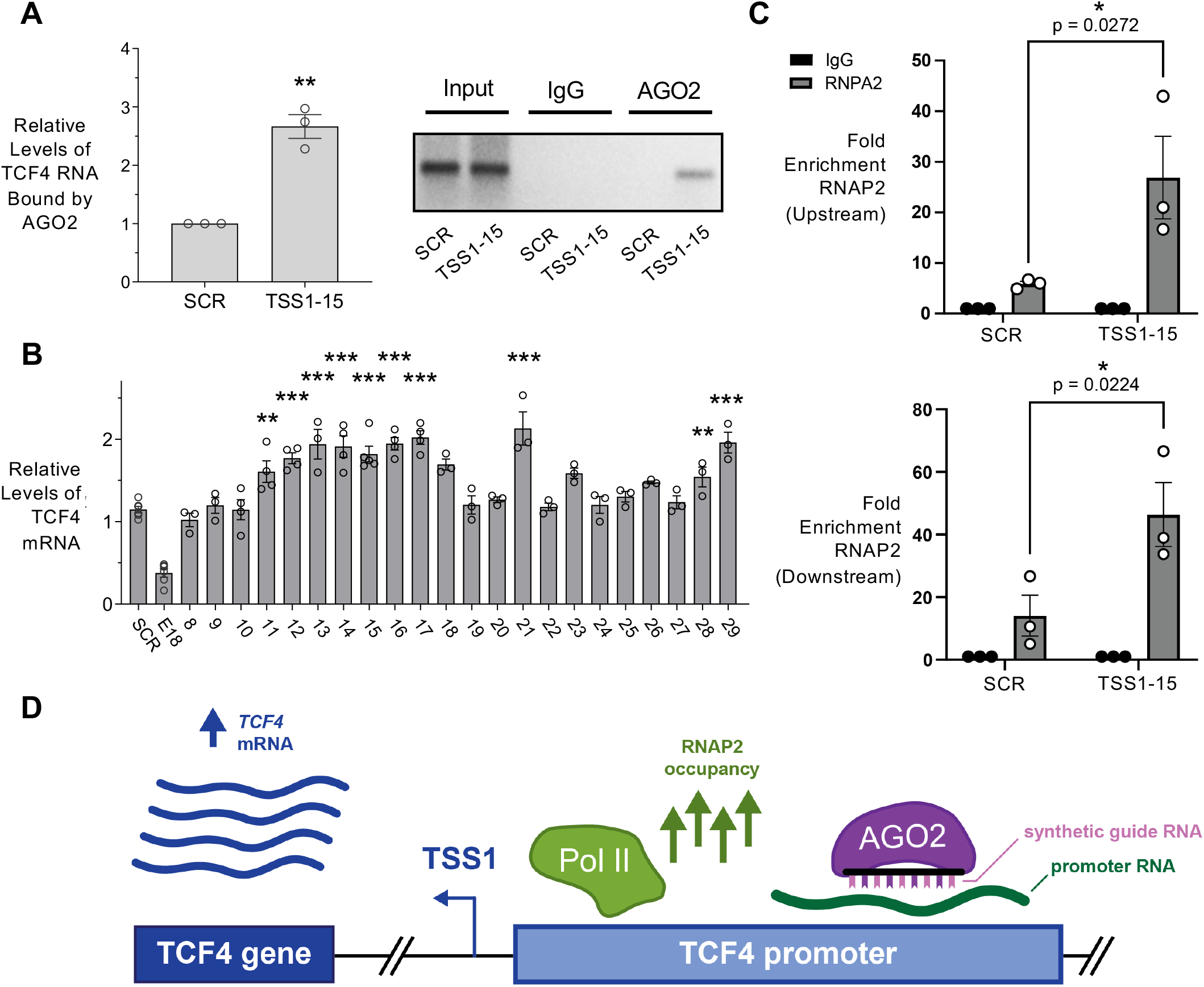
AGO2 association with *TCF4* promoter RNA and increased RNAP2 occupancy. **(*A*)** Nuclear RNA immunoprecipitation demonstrating recruitment of AGO2 to promoter RNA at the *TCF4* locus upstream from the TSS1 site upon addition of TSS1-15, relative to the SCR control. **(*B*)** Effects of dsRNAs TSS1-8 through TSS1-29 on *TCF4* mRNA expression analyzed by qPCR. Numbers are “fold change” relative to the scrambled dsRNA SCR. qPCR primers targeting *TCF4* exon 18 were used for quantification of *TCF4* mRNA levels. Sample significance for (A) and (B) was determined using a two-tailed Welch’s t-test. Statistical significance is indicated as follows: ∗p ≤ 0.05, ∗∗p ≤ 0.01, and ∗∗∗p ≤ 0.001. **(*C*)** RNAP2-ChIP-qPCR data demonstrating recruitment of RNAP2 at the TSS1 promoter region, using primer sets to detect association of RNAP2 both immediately upstream and downstream. Statistical significance (p-value ≤ 0.05) was determined using two-way analysis of variance and Tukey’s post-hoc test. **(*D*)** Scheme showing mechanism for RNA-mediated activation of gene expression that is supported by our data and consistent with previously published studies (Matsui et al. 2013; Johnson and Corey 2023). All data are from ≥ 3 independent biological replicates.

### TSS1-15 increases RNA polymerase II occupancy at the *TCF4* promoter

To test whether dsRNA-mediated TCF4 activation might be associated with enhanced transcription, we examined the expression of *TCF4* RNA by quantitative PCR (qPCR) and RNAP2 occupancy at the TSS1 promoter region by chromatin immunoprecipitation (ChIP). We observed increased expression of *TCF4* RNA by several promoter-targeted dsRNAs, including TSS1-15 (**Figure 4B**). Relative to the scrambled control, TSS1-15 increased RNAP2 occupancy both upstream and downstream of TSS1 (**Figure 4C**). As a positive control for the ChIP assay (Hofman et al. 2025), we also assessed RNAP2 occupancy at the promoter region of GAPDH (**Supplemental Figure S4B**). Our results, from both RNA immunoprecipitation and RNAP2 ChIP, support a model in which promoter-targeted dsRNAs act through AGO-associated recognition of promoter-region RNA, increase RNAP2 occupancy, and enhance transcription of *TCF4* (**Figure 4D**).

## DISCUSSION

### A hotspot for RNA-mediated activation of TCF4

In this study, we identify promoter-targeted dsRNAs that increase *TCF4* RNA and protein expression. Several active dsRNAs cluster within a defined region adjacent to the principal *TCF4* transcription start site, identifying a promoter hotspot that supports productive activation. Activity is retained by a dsRNA containing extensive 2′-fluoro, 2′-O-methyl, and phosphorothioate modifications commonly incorporated into therapeutic oligonucleotides. Anti-AGO2 immunoprecipitation enriches RNA transcribed from the targeted promoter region, and activation is associated with increased RNAP2 occupancy upstream and downstream of the principal transcription start site. These findings support a model in which AGO-associated recognition of promoter-region RNA enhances transcription of *TCF4*.

### Haploinsufficient disease and gene activation

Haploinsufficient disorders present a distinct therapeutic challenge because disease results from inadequate expression of a functional protein (Johnson et al. 2019; Evans et al. 2024; Tse and Corey 2026). Treatments directed at downstream symptoms may provide important clinical benefits but do not correct the underlying protein deficiency. Splice-switching ASOs offer one strategy for increasing protein expression when a suitable endogenous splicing event is targetable (Hofman and Corey 2024). Nusinersen increases productive *SMN2* expression in spinal muscular atrophy (Mercuri et al. 2018), and exon-skipping ASOs promote production of internally deleted dystrophin in Duchenne muscular dystrophy (Frank et al. 2020). Zorevunersen, an investigational ASO for Dravet syndrome, similarly increases productive expression from the functional *SCN1A* allele by modulating a non-productive splicing event (Laux et al. 2026). These examples demonstrate the therapeutic potential of increasing endogenous protein production, but their mechanisms depend on gene-specific opportunities to manipulate RNA processing.

Promoter-targeted dsRNAs offer a complementary approach that does not require a favorable splicing event (Johnson and Corey 2023). Because sequence recognition can be programmed by changing the guide strand RNA, the strategy could potentially be adapted to many different promoter regions. It may be particularly useful for genetically heterogeneous disorders in which many distinct loss-of-function mutations reduce expression from a single allele. Rather than correcting each pathogenic variant individually, increasing transcription from the remaining functional allele could provide a mutation-independent approach applicable to many patients. We note, however, that our approach is not intrinsically selective for the functional allele. It may also activate expression of the mutant allele, and the downstream effects of any harmful products should be considered.

### General lessons for RNA-mediated activation

In most of our previous studies of RNA-mediated gene activation, we did not prioritize the disease relevance of the target. We mostly targeted genes that had well characterized promoters and well understood basal and activated states. One exception was the *MECP2* gene. *MECP2* is implicated in Rett syndrome, with the wild-type copy silenced by DNA methylation. Activation might provide therapeutic benefit. In a clonal cell line with both alleles silenced, we observed no detectable RNA transcription at the gene promoter and no evidence for increased gene expression after addition of promoter-targeted dsRNAs (Yu et al. 2011). The absence of expression of promoter RNA is consistent with our inability to observe gene activation. Our experience with *MECP2* is a reminder that target genes with high unmet need for therapeutic upregulation may lack the mechanistic foundation necessary for RNA-mediated activation.

In this case, the haploinsufficiency of *TCF4* and the need for a single agent to treat most patients create high therapeutic need. The *TCF4* gene, while not as challenging as *MECP2*, presented difficult obstacles to the design of candidate RNAs. The promoter is TATA-less with a complex architecture and several CpG islands. The RNA landscape at the gene is not well characterized and it has at least three putative promoters. Transcription from these promoters and alternative splicing generate multiple TCF4 transcript variants.

Another challenge presented by activation of *TCF4*, which will be shared by other attempts to activate gene expression for haploinsufficient disease, is that the level of activation that would restore normal abundance is ∼2-fold. In our previous studies RNA-mediated activation had been as high as 20 to 30-fold (Janowski et al. 2007; Matsui et al. 2013), making the observation of gene activation simple. The biologically-relevant levels of activation that we observed required an unusually high number of replicates and the development of rigorous protocols for minimizing the noise of expression assays.

We find that these challenges can be overcome by close examination of existing databases to lay a foundation for understanding transcription at gene promoters. This computational analysis is then complemented by experimental measurement of transcription to confirm the RNA transcripts that provide potential binding locations for RNA-mediated control. Finally, testing multiple RNAs allows the development of rigorous structure-activity relationships that enable an understanding of targeted gene activation.

#### Activating RNAs cluster around a target hotspot

The clustering of multiple active dsRNAs at the TSS1 region suggests that promoter activation depends on more than simple complementarity to any transcript near the promoter. Neighboring guide sequences may position AGO-containing complexes within a local promoter-RNA environment that is permissive for transcriptional activation. The identification of active, overlapping dsRNAs provides starting points for optimization and suggests that productive engagement is tolerated across a limited sequence window. Anti-AGO2 immunoprecipitation supports association of AGO2 with RNA from this region, although these experiments do not by themselves establish direct base pairing between the synthetic guide and promoter RNA or a direct physical interaction between AGO2 and RNAP2.

Retention of activity after extensive chemical modification is important for therapeutic development. Unmodified duplex RNAs have limited metabolic stability, and modifications that increase stability can alter AGO loading, guide-strand selection, and target recognition (Egli and Manoharan 2023; Hofman and Corey 2024). The activity of Mod15 therefore demonstrates that promoter-targeted activation is compatible with a chemical architecture resembling those used in therapeutic siRNAs. Further optimization will be required to define the modification patterns, potency, duration of action, and selectivity most suitable for gene activation.

Advances in extrahepatic oligonucleotide delivery broaden the potential relevance of this strategy. Chemically modified and multivalent siRNA scaffolds can support durable activity in the central nervous system (Alterman et al. 2019; Brown et al. 2022), while ligand- and antibody-based conjugates are extending functional oligonucleotide delivery to additional tissues, including skeletal muscle (Malecova et al. 2023). These delivery platforms were developed primarily for gene silencing, and their compatibility with promoter-targeted activation remains to be established. Nevertheless, they provide a growing technological foundation for evaluating activating dsRNAs in tissues relevant to haploinsufficient disease.

### Conclusions

In this study, we report that RNA-mediated gene activation can be applied to increasing expression of the *TCF4* gene responsible for Pitt Hopkins Syndrome. The approximately 2.0 to 3.0-fold activation observed is comparable in magnitude to the increase that might compensate for reduced dosage due to the mutant allele. We identify promoter-targeted dsRNAs that increase endogenous *TCF4* expression, define a sequence-dependent hotspot for productive activation, and retain activity after extensive chemical modification. Our data provide chemically tractable starting points for developing *TCF4*-activating compounds. While further studies in patient-derived iPSC-derived cells will be needed to advance compounds tor Pitt Hopkins Syndrome, our data support broader investigation of promoter-targeted RNAs as a mutation-independent strategy for haploinsufficient disease and other diseases that would benefit from modest activation of gene expression.

## MATERIALS AND METHODS

### Cell culture

A549 cells were obtained through the American Type Culture Collection (ATCC). A549 cells were cultured in Dulbecco’s Modified Eagle’s medium (DMEM) with high glucose (Millipore Sigma), supplemented with 10 % FBS. All cell cultures were grown at 37°C in 5 % CO2 and passaged when 70 %–80 % confluent.

### TCF4 DNA and RNA landscape analysis

All computational datasets used in this study to characterize the DNA and RNA landscape of *TCF4* were generated and published by major consortium efforts. Additional computational tools used in this study have been peer-reviewed. All datasets and tools applied in this study were readily available for public use in the UCSC Genome Browser (GRCh38/hg38) without restrictions.

### Standard RT-qPCR assay

Total RNA was extracted from cells using TRIzol reagent. Reverse transcription was performed using high-capacity reverse transcription kit (Applied Biosystems) per the manufacturer’s protocol. 2.0 μg of total RNA was used per 20 μL of reaction mixture. All qPCR reactions were set-up in 96-well optical plates using iTaq™ Universal SYBR® Green Supermix (Bio-Rad Laboratories), and were run on a CFX96 Touch Real-Time PCR Detection System (Bio-Rad Laboratories). All target-specific qPCR data were normalized to the relative levels of an internal control gene, *RPL19*. Target-specific primers were synthesized by Integrated DNA Technologies and are listed in Supplemental Table S1.

### Strand-specific RT-qPCR assay

To amplify and measure sense and antisense RNA species across the *TCF4* promoter region, we adapted strategies from prior work by our laboratory (Matsui et al. 2013). Briefly, primer sets were designed to amplify regions between TSS3 and the first coding exon shared between all prioritized *TCF4* transcripts identified by our GENCODE analysis. Specific primers were also designed to generate sense and antisense qPCR templates to distinguish strand-specific RNA species at the targeted regions. These primers were synthesized by Integrated DNA Technologies and can be found in Supplemental Table S1.

Total RNA was purified from cultured A549 cells using phenol–chloroform extraction and ethanol precipitation, followed by DNA digestion using DNase I (NEB). DNase-treated total RNA was used for strand-specific reverse transcription using the high-capacity cDNA reverse transcription kit (Applied Biosystems). Primer TCF4-23R was used to generate cDNA from sense RNA, whereas AS-TCF4 was used to generate cDNA from antisense RNA. Thirty nanograms of either sense or antisense cDNA was used as a qPCR template for the iTaq™ Universal SYBR® Green Supermix reaction following the manufacturer’s instructions (Bio-Rad Laboratories). All qPCR reactions were performed in 384-well optical plates and were run on a CFX384 Touch Real-Time PCR Detection System (Bio-Rad Laboratories). All target-specific qPCR data were normalized to the relative levels of an internal control gene, *RPL19*.

### Synthesis and formation of dsRNAs

Sense and antisense strands for each unmodified or chemically modified dsRNA tested in this study targeting *TCF4* were synthesized by Integrated DNA Technologies. For promoter-targeted dsRNAs, each sense and antisense strand contains 19 ribonucleotides specific to its target along with two non-specific deoxythymidine overhangs at the 3’ end, comprising a 21-mer dsRNA. The design of the siRNA specific to *TCF4* was synthesized as previously described as a 27-mer (Forrest et al. 2013). Respective sense and antisense strands are then annealed together to form dsRNA in the same solution by denaturing at 95°C and allowing it to gradually cool down to room temperature in a thermocycler. The nucleic acid sequence for each sense and antisense strand of a given unmodified dsRNA from this study can be found in Supplemental Table S2.

### Transfection of dsRNAs into cell cultures

All transfections used Lipofectamine RNAiMAX (Invitrogen) and reduced serum media, Opti-MEM™. Cells were seeded into six-well plates 24-hours before transfection, at 200,000 cells per well. The next day, a given dsRNA was transfected into the cells at a final concentration of 25 nM. 24-hours later, the transfection media was replaced with full culture media. Cells were then harvested for western analyses 72-hours after transfections. Transfections were designed using standard guidelines for rigor and controls (Gagnon and Corey 2019).

### Western blotting analysis

Proteins were separated on 4%–20% gradient Mini-PROTEAN® TGX precast gels (Bio-Rad) using a Mini-PROTEAN Tetra Vertical Electrophoresis Cell (Bio-Rad). After gel electrophoresis, proteins were semi-dry transferred (Power Blotter, Invitrogen) to a nitrocellulose membrane (0.45 μm, Cytiva) using the “Mixed Range MW” setting (25V, constant 1.3A, and 7 minutes) and Extra Thick Western Blotting Paper (Thermo Fisher). Membranes were blocked for 1 h at room temperature with 5% milk in 1× PBS containing a final 0.05% of Tween-20 (v/v) (PBST). Blocked membranes were incubated with the following primary antibodies at 4°C on a rocking platform overnight: anti-TCF4 1:1000 (Santa Cruz) or anti-β-actin 1:10000 (Sigma-Aldrich). After primary antibody incubation, membranes were washed 3× for 10 min at room temperature with PBST and then incubated for 1 hr (anti-TCF4) or 30 min (anti-β-actin) at room temperature with respective secondary antibodies in blocking buffer. Membranes were washed again 3× 10 min in PBST. Washed membranes were soaked with horseradish peroxidase chemiluminescence substrate according to manufacturer’s recommendations (SuperSignal™ West Pico Chemiluminescent substrate, Thermo Scientific, Clarity™ Western ECL substrate, Bio-Rad, or Clarity™ Max Western ECL substrate, Bio-Rad) and imaged. The membranes were imaged using the ChemiDoc ™ Go Imaging system (BioRad), and the bands were quantified using Image Lab 6.1 (Bio-Rad). Protein quantification in Image Lab was performed using default settings and the instructions provided by the manufacturer (Bio-Rad).

### RNA immunoprecipitation assay (RIP)

3.0 million A549 cells were seeded in a 150 mm dish the day before the cationic transfection of dsRNAs (2 dishes per dsRNA treatment). The media in each cell culture was replaced with fresh DMEM 24-hours post-transfections. Cells were harvested 48-hours post-transfection. Detached cells were fractionated according to our previous protocol (Hofman et al. 2025). The separated cell nuclei were lysed in whole cell lysis buffer (20 mM Tris-HCl pH7.4, 150 mM NaCl, 1 mM EDTA, 0.5% NP-40, 0.5 mM DTT, protease inhibitor, and RNase inhibitor (Promega, 50 U/ml final)) with a volume 3 times the volume of the cell pellet size. After mixing thoroughly, the lysate was stored at -80°C overnight. Insoluble material was removed by centrifugation at 12,000×g for 15 min at 4°C. The supernatant was isolated and quickly frozen in liquid nitrogen and stored.

60 µL Protein A/G agarose Plus beads were washed with 1×PBS (pH 7.4) and incubated with 5 µl of anti-AGO2 antibody (Wako) or mouse IgG in 1×PBS (pH 7.4) at 4°C with gentle agitation for 2 hours. After two washes of 1×PBS, beads were incubated with isolated cell lysate for 2hr at 4 °C. The beads were extensively washed with 1×PBS (pH 7.4) once, washed twice with IP wash buffer (150 mM NaCl, 3 mM MgCl_2_, 0.5% NP-40, and 20 mM Tris-HCl (pH 7.4)), and finally once more with 1×PBS (pH 7.4). The beads were finally eluted with elution buffer (1 % SDS, 0.1 M NaHCO_3_ and RNase inhibitor). After proteinase K treatment, RNA extraction and precipitation, samples were treated with recombinant DNase I, followed by reverse transcription. Corresponding cDNA was amplified in a standard RT-qPCR assay using primers complementary to regions upstream of the *TCF4* transcription start site (see Supplemental Table S1). The PCR products were analyzed using a 3 % agarose gel, and the expected amplicon size was predicted *in silico* and confirmed visually when compared to an appropriate DNA size standard.

### RNAP2 ChIP-qPCR assay

2.5 million A549 cells were seeded in a 150 mm dish the day before the cationic transfection of dsRNAs. The media in each cell culture was replaced with fresh DMEM 24-hours post-transfections. 48-hours post-transfection, cells were crosslinked with 1.0% formaldehyde at room temperature for 10 min, followed by the addition of glycine (final concentration of 150 mM) to quench the crosslinking reaction at room temperature for 5 minutes. Crosslinked cells were harvested using a cell scraper, transferred into a conical tube, and pelleted by centrifugation (500 rcf) at 4°C. Nuclei from crosslinked cells were then isolated with hypotonic lysis buffer (10 mM Tris–HCl pH 7.4, 10 mM NaCl, 3 mM MgCl_2_, 2.5% NP-40, 0.5 mM DTT, protease inhibitor) and lysed in lysis buffer (50 mM Tris–HCl pH 7.4, 120 mM NaCl, 0.05% NP-40, 1 mM EDTA, 1 mM DTT, protease inhibitor). These nuclear lysates were then fragmented by sonication for 20 minutes (30 s on, 30 s off, at 50 % amplitude using a Qsonica Q800R3). A fraction of the fragmented nuclear lysate (5.0 % of total eluate recovered) was used to evaluate crosslinking and fragmentation quality prior to performing immunoprecipitation.

As previously performed in our laboratory (Hofman et al. 2025), nuclear lysate was precleared with 1 µg of normal mouse IgG (12-371, Millipore Sigma) using protein G Dynabeads (Invitrogen), rotating for 1 hour at 4°C. Precleared nuclear lysate was then isolated and incubated with 3 µg of either the RNAP2 antibody (05-623, Millipore Sigma) or the normal mouse IgG in immunoprecipitation buffer, rotating overnight at 4°C. Antibody–protein–DNA complexes were then recovered with protein G Dynabeads, rotating for 3 hours at 4°C. Captured complexes on the beads were washed with low salt, high salt, LiCl, and TE buffers. Each wash step was performed with rotation for 5 minutes at 4°C. After washes, captured complexes were eluted twice with 250 μL of elution buffer (1% SDS, 0.1 M NaHCO_3_) at room temperature. Crosslink reversal of immunoprecipitated protein-DNA complexes was achieved by adding NaCl (final concentration of 200 mM) and incubation at 65°C for 2 hours. Proteins in the immunoprecipitated samples were digested using proteinase K and heat at 42°C for 50 min. DNA from immunoprecipitated samples was then recovered using phenol–chloroform extraction and ethanol precipitation. ChIP DNA samples were dissolved in nuclease-free water and analyzed by qPCR. Primer sets used for RNAP2-ChIP are listed and indicated in Supplemental Table S1.

### Statistical analyses

Each experimental replicate was produced by a separate transfection experiment, usually on different days and often by a different experimenter. Each sample was analyzed by western analysis multiple times, and these technical replicates were used to produce the reported biological value.

Quantification of total TCF4 protein and mRNA levels from western blots, standard qPCR, and AGO2-RIP-qPCR was performed using Excel and GraphPad Prism 11. All data are expressed as mean ± the standard error of the mean (SEM) and represent the results of at least three independent biological replicates (Supplemental Figure S3), as reflected by each individual datapoint within the bar graphs. “n” represents the number of biologically independent samples, as specified in the corresponding figure legends. Technical replicates were averaged into respective biological replicates. For western analysis, raw values corresponding to TCF4 were normalized to the raw values of its corresponding loading control, β-actin. Fold change between conditions was calculated by dividing the normalized values of an experimental condition by the normalized values of the appropriate control condition. A quantile-quantile plot and a Brown-Forsythe test were performed to determine the dataset fit the assumptions made by Welch’s t-test. Sample significance was then determined using a two-tailed Welch’s t-test. Statistical significance is indicated as follows: ∗p < 0.05, ∗∗p < 0.01, and ∗∗∗p < 0.001.

Quantification of RNAP2 fold-enrichment between dsRNA conditions tested was performed in GraphPad Prism 11.0.0. All data are expressed as mean ± the standard error of the mean (SEM) and represent the results of three independent biological samples, as reflected by each individual datapoint within the bar graphs. “n” represents the number of biologically independent samples, as specified in the corresponding figure legends. Statistical significance was determined using an ordinary two-way ANOVA with a post-hoc Tukey’s multiple comparisons test. Statistical significance is indicated as follows: ∗p < 0.05, ∗∗p < 0.01, and ∗∗∗p < 0.001. Before data analysis, the experimental blot results were excluded from further analysis if the prespecified positive control E18 failed to reduce full length TCF4 by approximately 80% or if the TCF4 or beta-actin signal for any required control fell outside the validated range for quantitation. No individual lanes were selectively excluded. The SCR and E18 controls were used in every experiment.

## SUPPLEMENTAL INFORMATION

Supplementary material is available for this article.

Document S1. Supplemental Figures S1–S4

Document S2. Supplemental Tables S1-S2

## ACKNOWLEDGMENTS

D.R.C. was supported by the National Institutes of Health (NIH) (R35GM118103) and the Robert Welch Foundation (I-2184) and holds the Rusty Kelley Professorship in Medical Science. We acknowledge with gratitude support provided by the Vincent Pauca “Be Better” Memorial Award from the Pitt Hopkins Research Foundation and Audrey Davidow for inspiring us to initiate this project. Bethany Janowski was supported by a Gilman “Special Opportunities in Pharmacology Award” from the UT Southwestern Department of Pharmacology.

## AUTHOR CONTRIBUTIONS

D.R.C. and B.A.J. conceptualized the study. B.A.J., V.T., K.D., J.H., and S.A.C. performed all experiments. D.R.C. and V.T. wrote the manuscript with input from all authors. D.R.C. and B.A.J. supervised the work and were responsible for funding acquisition.

## AI Disclosure

The authors used ChatGPT (OpenAI) to improve the readability and proofread the text.

## Data availability Statement

Primary data is available upon request from the authors.

---

**Supplemental Figure S1.**
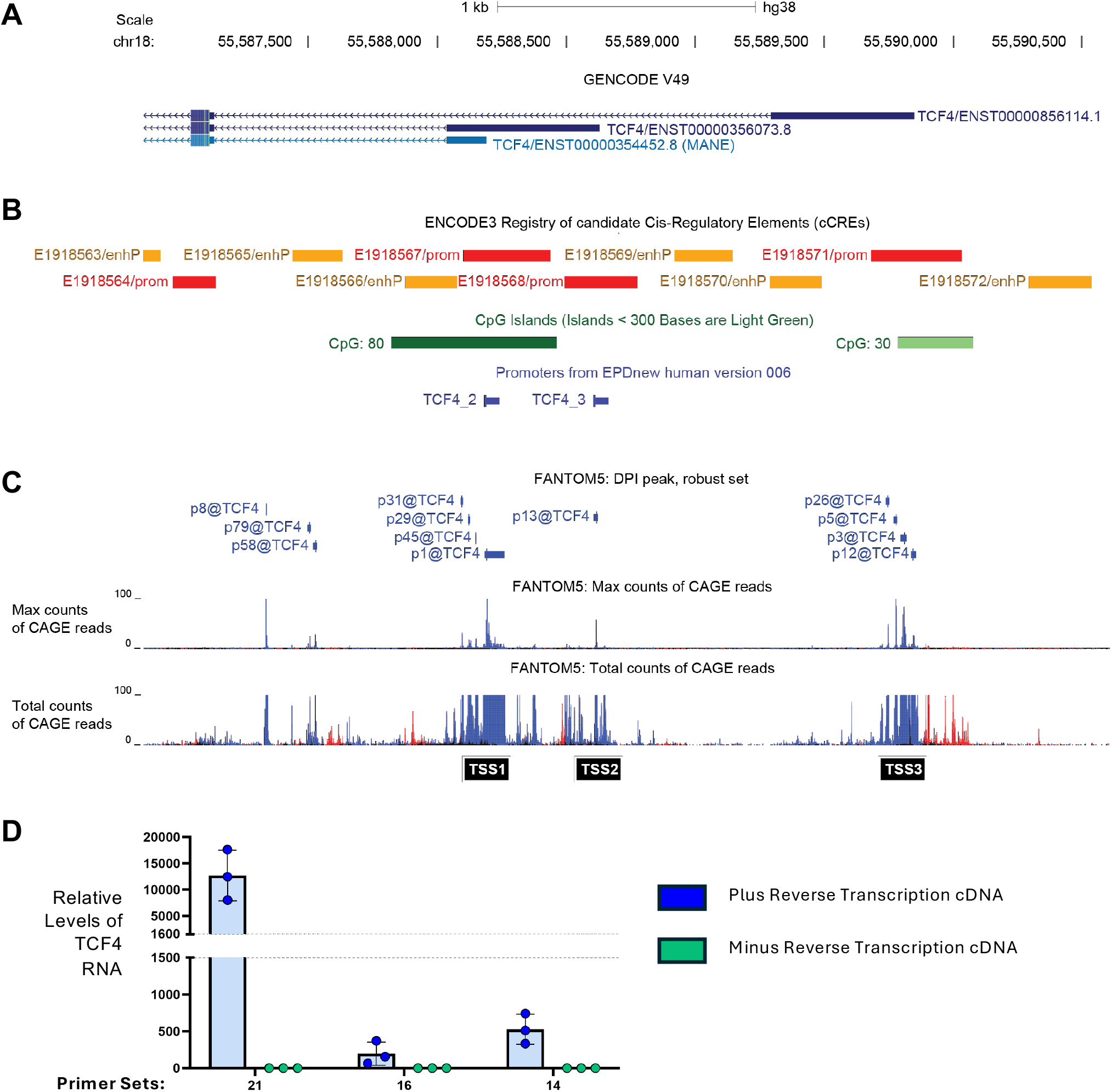
Transcription at the *TCF4* promoter. **(*A*)** Prioritized GENCODE transcripts are shown. The MANE *TCF4* transcript variant is shown in teal. Non-MANE transcripts are shown in dark blue. Thick and thinner boxes represent exons and untranslated regions, respectively. Lines with arrows represent introns and the directionality of the sense strand. **(*B*)** The ENCODE track maps where promoter-like (red) or promoter-enhancing (orange) elements are relative to the prioritized *TCF4* transcripts. The CpG island analysis track identifies CG dinucleotide repeats within the region of interest, with one containing 80 CG dinucleotides (dark green) and another containing 30 CG dinucleotides (light green). The EPDnew track shows predictions for two putative promoters within the region of interest (shown in blue). **(C)** CAGE-sequencing dataset by the FANTOM5 consortium is shown. The total coverage of CAGE-seq reads (bottom), and the max count and clustering of CAGE-seq reads (top), are shown for the 5’ end of the TCF4 locus. Sense strand transcription is annotated in blue. Antisense strand transcription is indicated in red. The annotation of TSSs (supported by ENCODE, CAGE, and *in silico* tools) are indicated at the bottom of the figure in black and white text as TSS1, TSS2, and TSS3. ***(D)*** qPCR (linear scale) showing detection and amplification of promoter-specific RNA at the TSS1 region is RNA dependent and not a result of contaminating and/or genomic DNA. Data are from ≥ 3 independent biological replicates.

**Supplemental Figure S2.**
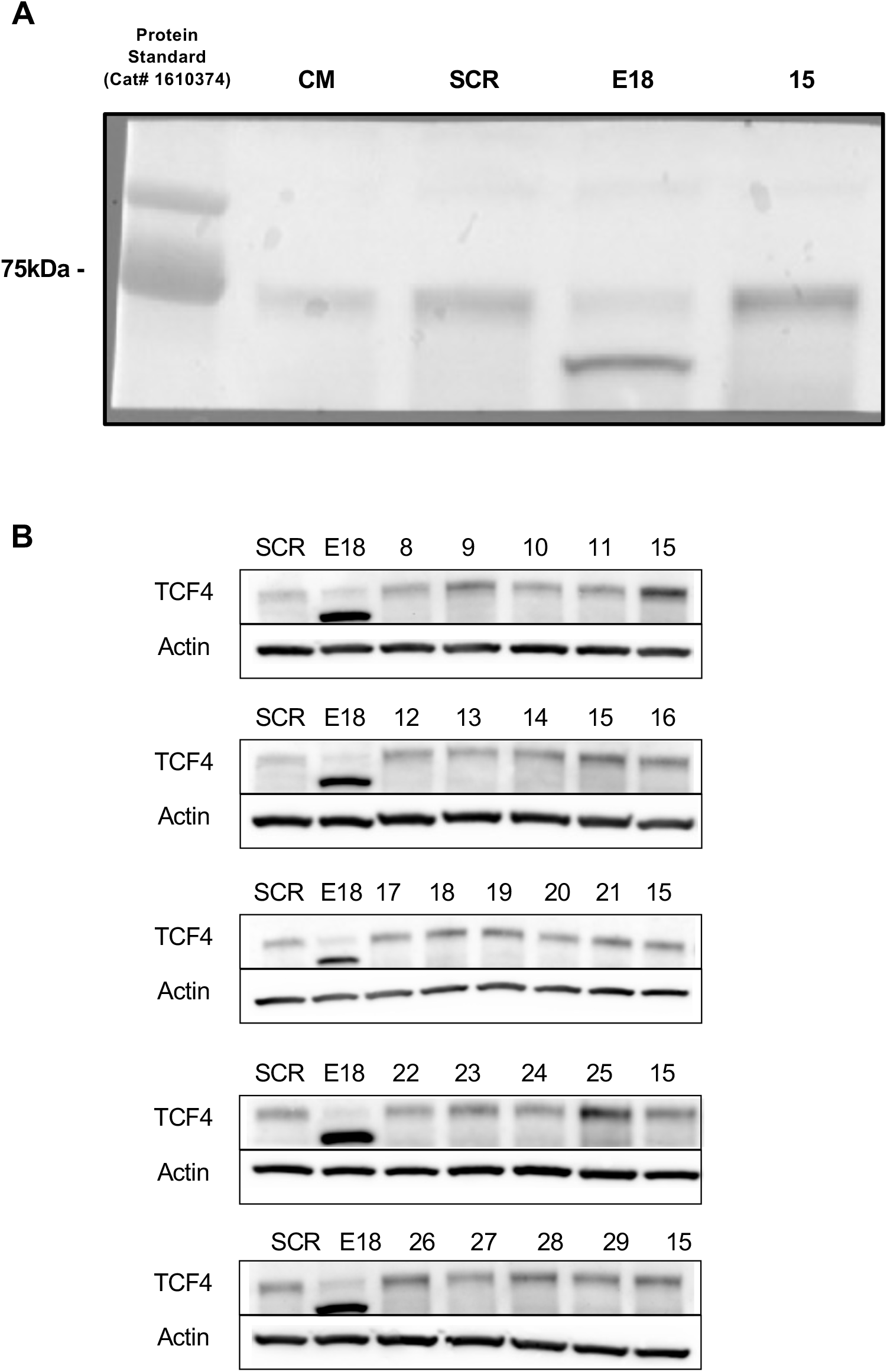
Uncropped and cropped representative western blots for lead activator TSS1-15 and the tiled dsRNA screen. (*A*) Uncropped blot corresponding to the representative western blot shown in Figure 2B. A protein size standard is shown in the first lane. (*B*) Cropped, representative western blots for the tiled dsRNA screen corresponding to Figure 2D.

**Supplemental Figure S3.**
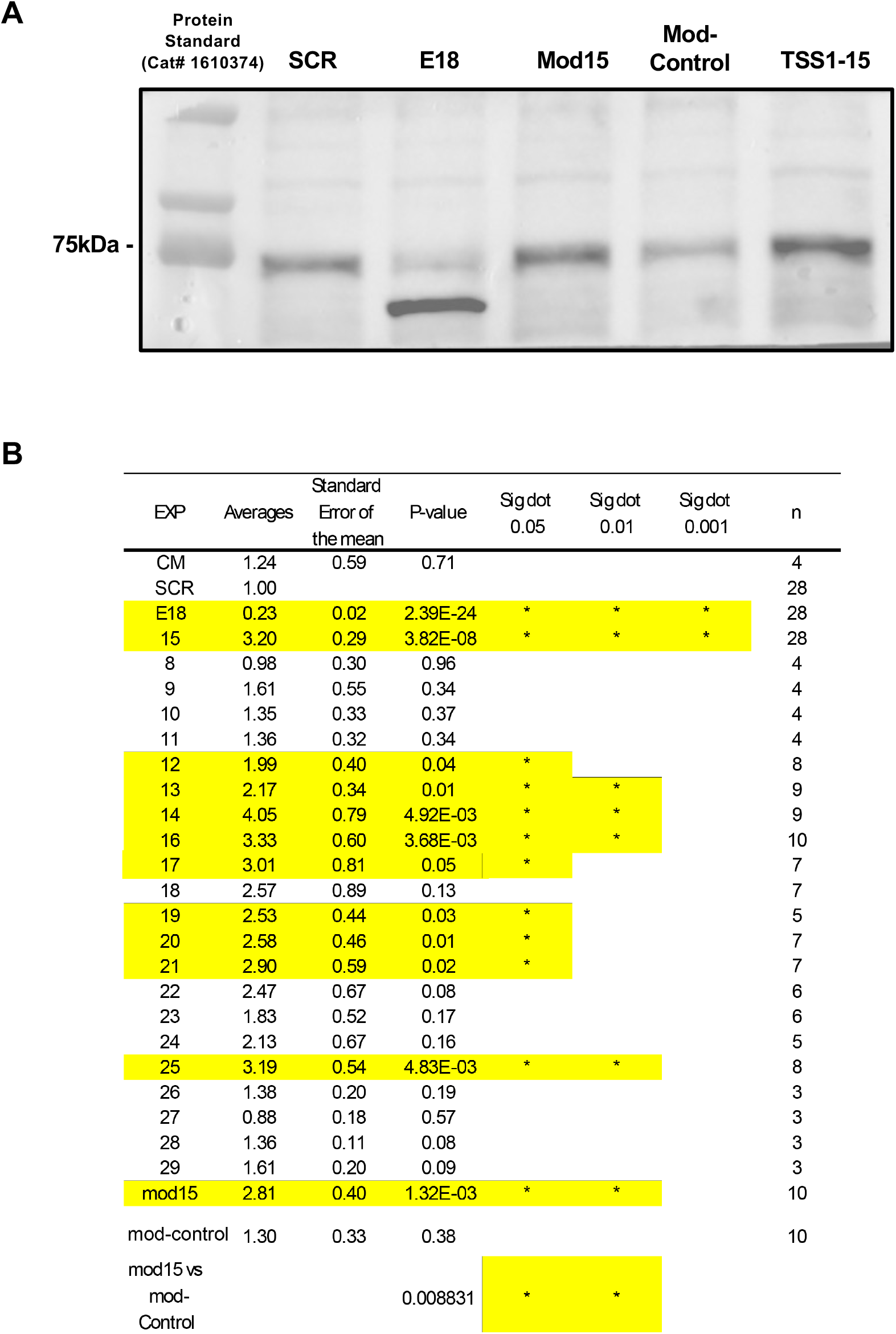
Uncropped representative western blots comparing lead activator TSS1-15 to a chemically-modified analogue. (*A*) Uncropped blot corresponding to the representative western blot shown in Figure 3B. A protein size standard is shown in the first lane. (*B*) Summary of biological replicate numbers and p-values.

**Supplementary Figure S4.**
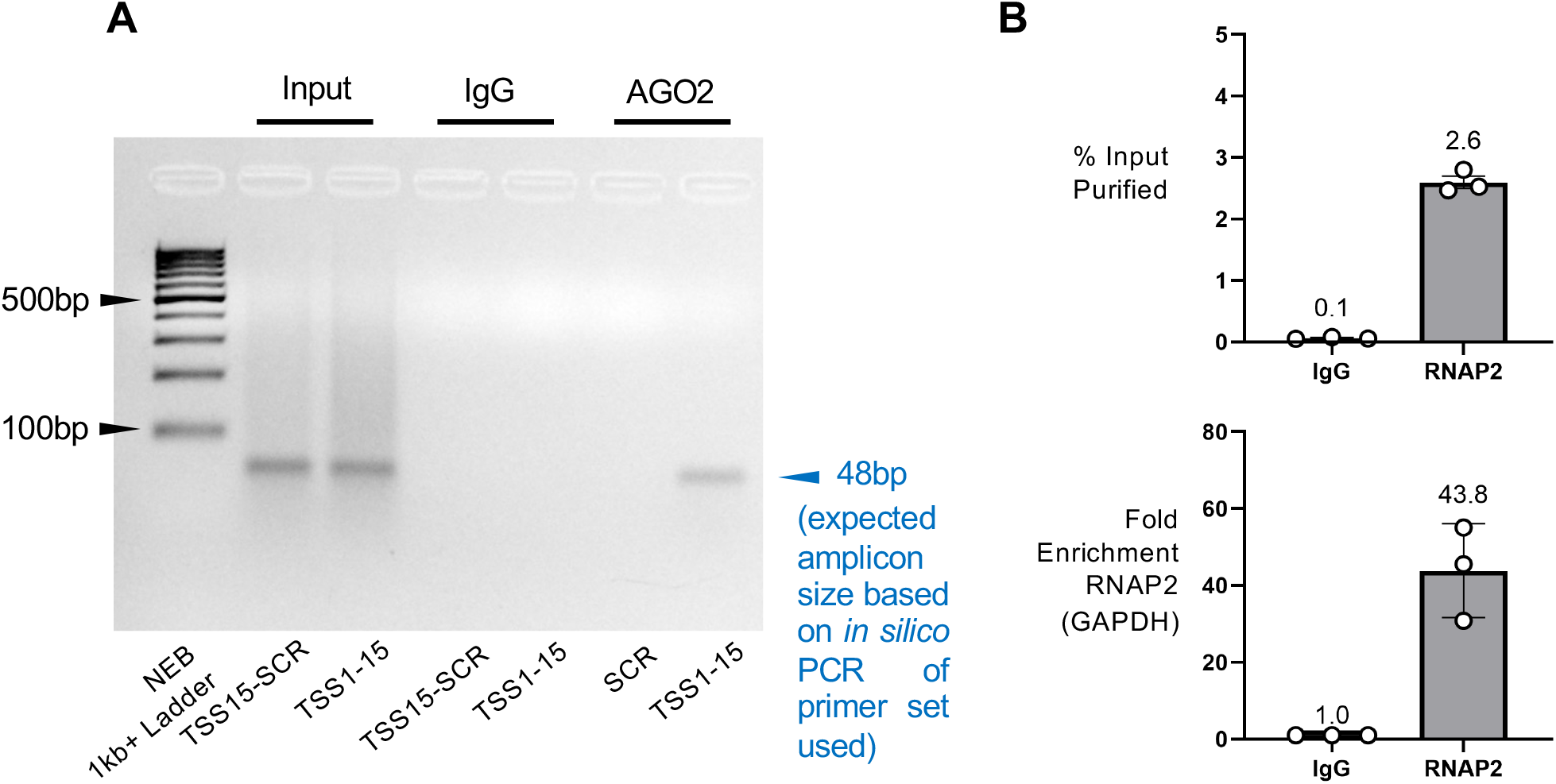
Association of TSS1-15 with promoter transcripts and its impact on *TCF4* RNA synthesis. (*A*) Uncropped agarose gel of RT-qPCR products corresponding to Figure 4A. A DNA size standard is shown in the first lane. The expected amplicon size across all conditions is indicated in blue text. (*B*) RNAP2-ChIP-qPCR targeting the promoter region of GAPDH, an internal control gene. Demonstrates successful chromatin immunoprecipitation in our hands (represented as percent input purified from nuclear lysate), and successful detection of enriched RNAP2 occupancy relative to a negative IP control (represented as fold-enrichment).

**Supplemental Tables S1.**
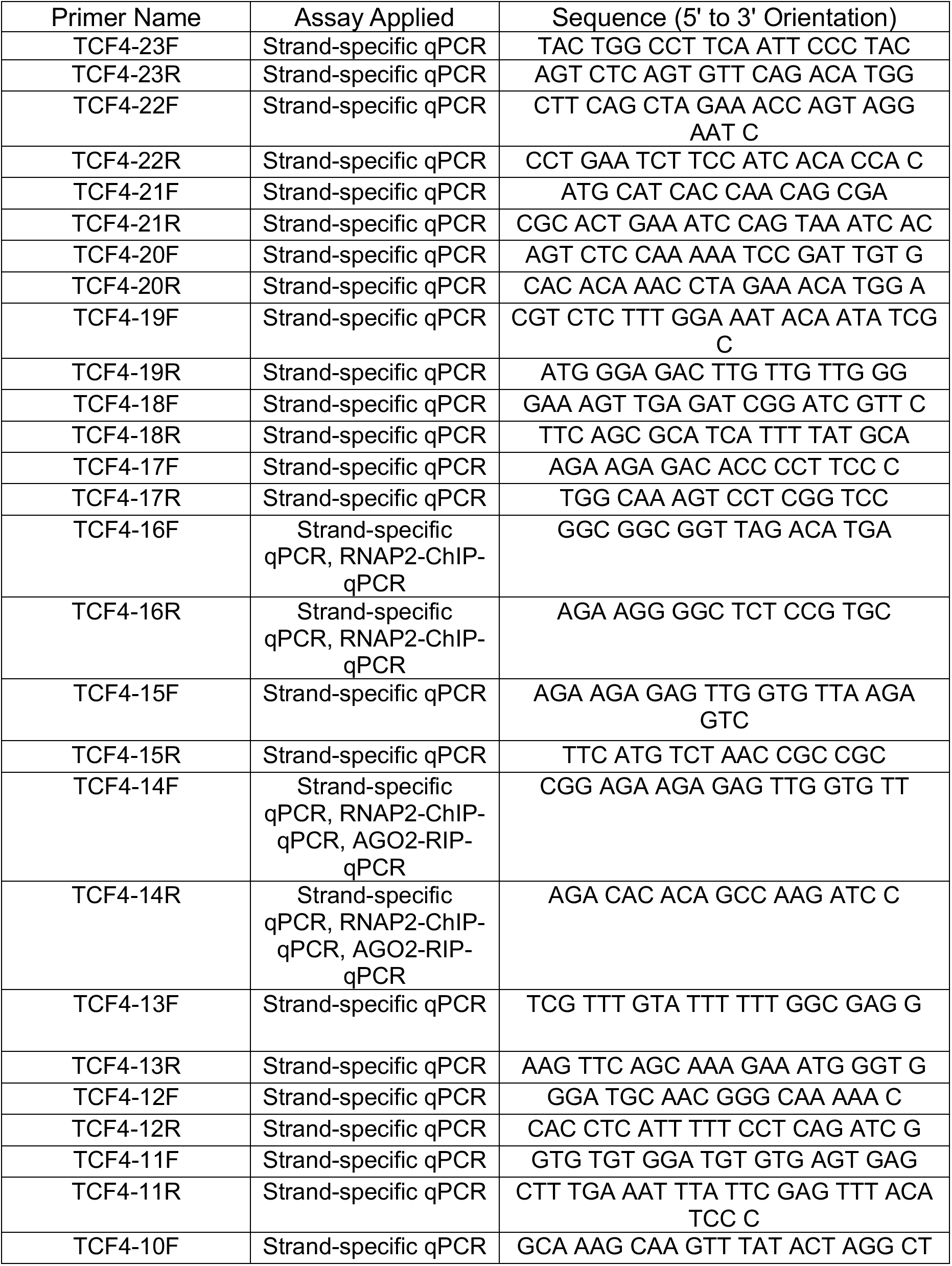
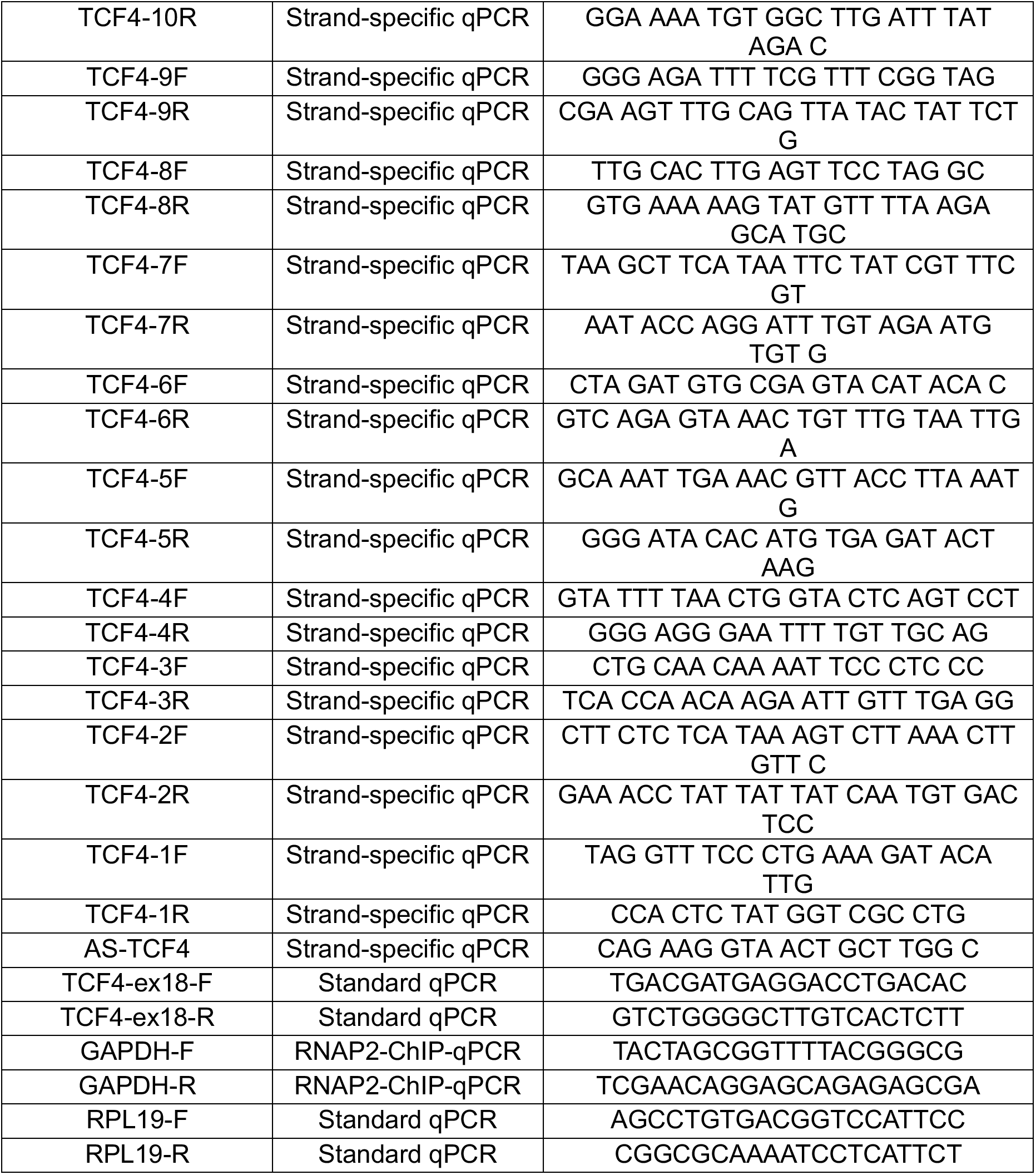
A list of primer sequences designed, validated, and applied for various qPCR assays presented in this study.

**Supplemental Tables S2.**
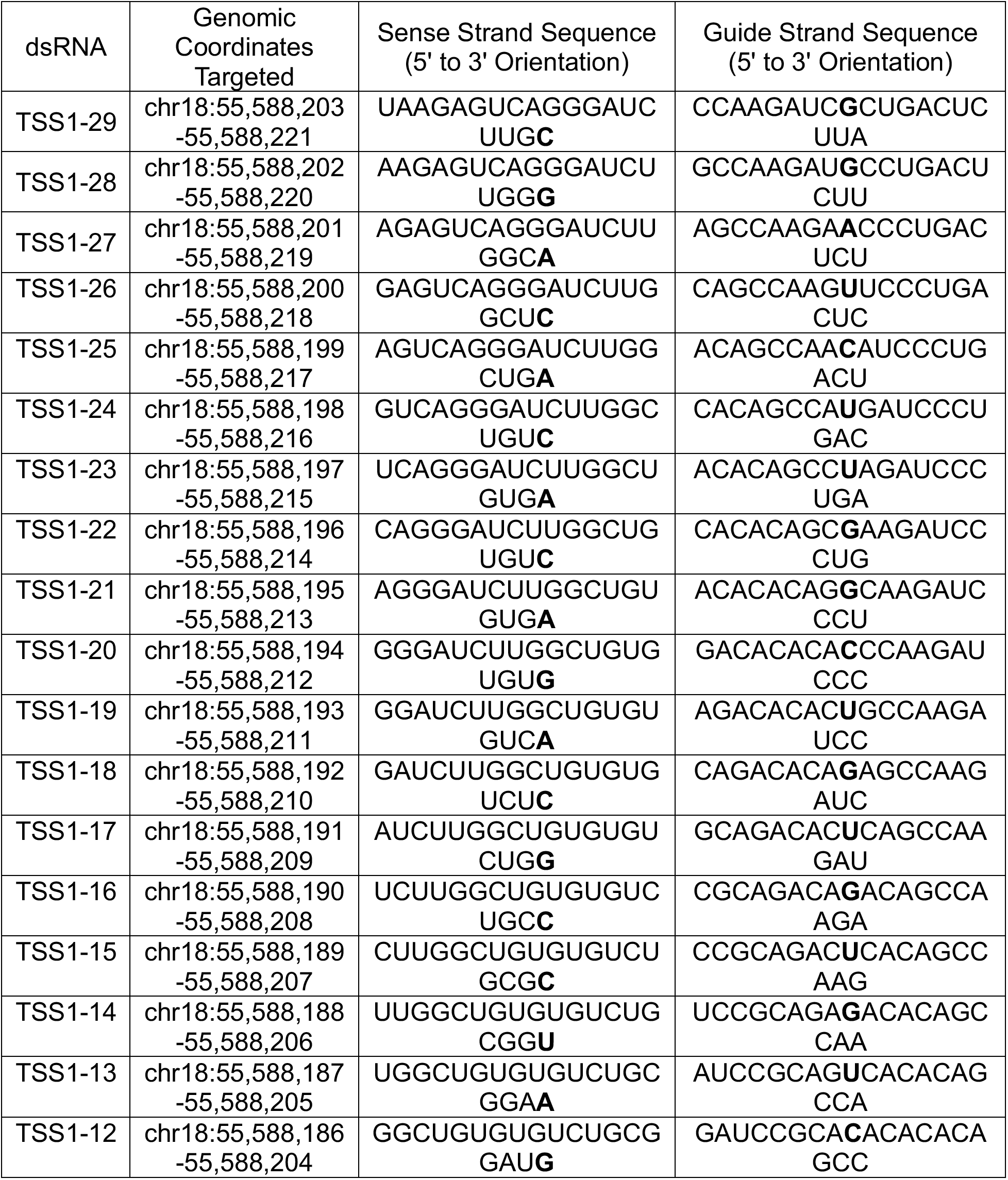
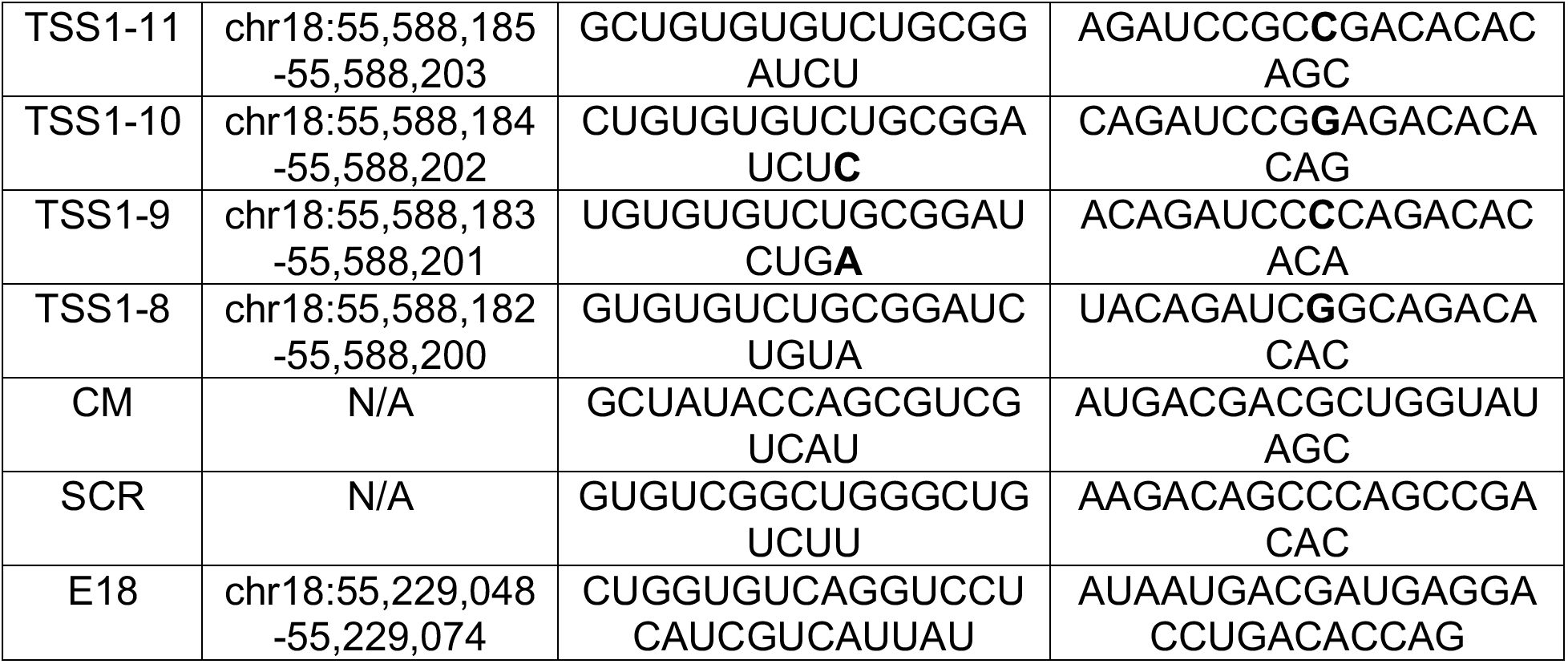
A list of dsRNAs designed and screened in this study. All sequences show ribonucleotides (double deoxythymidine overhangs at the 3’ end of each sense or guide strand sequence is not shown in the table below). Bolded text indicate positions of introduced mismatches.

